# How Cognitive and Affective Empathy Relate to Emotion Regulation: Divergent Patterns from Trait and Task-Based Measures

**DOI:** 10.1101/611301

**Authors:** Nicholas M. Thompson, Carien M. van Reekum, Bhismadev Chakrabarti

## Abstract

Evidence suggests that empathy and emotion regulation may be related, but few studies have directly investigated this relationship. Here we report two experiments which examined: 1) how different components of empathy (cognitive & affective) relate to the habitual use of cognitive reappraisal to regulate emotions (N=220), and 2) how these components of empathy relate to implicit reappraisal in a context-framing task (N=92). In study 1, a positive correlation between cognitive empathy and reappraisal use was observed. Affective empathy showed no relationship with reappraisal use. In study 2, participants completed an implicit reappraisal task in which previously viewed negative images were paired with either a neutralising (intended to reduce negative emotionality) or descriptive (which simply described the image) framing sentence. Participants then reported how unpleasant/pleasant each image made them feel. In contrast to study 1, a positive correlation between affective empathy and the implicit reappraisal task metric (rating of neutralising–descriptive framing conditions) was observed. There was no relationship between cognitive empathy and implicit reappraisal. These findings suggest that both components of empathy are related to reappraisal, but in different ways: Cognitive empathy is related to more deliberate use of reappraisal, while affective empathy is associated with more implicit reappraisal processes.

---

Empathy and emotion regulation represent distinct socio-emotional abilities, both of which are critical for effective social functioning and emotional wellbeing (Decety & Lamm, 2006; Fodor, 1987). Empathy refers to a set of processes involved in understanding and responding appropriately to the mental states and emotions of others (Chakrabarti & Baron-Cohen, 2006). Emotion regulation refers to the ability to exert control over one’s emotional experiences (Gross, 1998). Even though these processes are distinct, they often interact in everyday social behaviour: Consider a situation in which somebody has upset or angered you. Trying to see things from that person’s perspective and understand the motivations behind their actions can often help to regulate any negative emotions we felt as a result of their behaviours. Or think of a parent whose child is in pain. Through affective empathy the parent may share their child’s distress. However, if they are unable to sufficiently regulate this emotional state, such distress could impede the parent’s ability to comfort and support their child appropriately. These two everyday examples just scratch the surface of the complex interaction between empathy and emotion regulation. While extant work summarised below provides evidence to suggest that these abilities may be related, the nature of the inter-relationship between different component processes related to empathy and emotion regulation is not well characterised. In our research presented here, we examine the relationship between trait empathy and emotion regulation using a combination of trait and task measures.

Empathic abilities have been shown to vary within the general population and are considered to be measurable and relatively stable traits (Baron-Cohen & Wheelwright, 2004). Most theories of empathy distinguish between two core facets: affective empathy (the capacity to *feel* what another individual is feeling), and cognitive empathy (the capacity to *understand* the emotional state of another individual through the use of inferential cognitive processes) (Chakrabarti & Baron-Cohen, 2006; Singer & Lamm, 2009). Dual process models of empathy have garnered much support, and in alignment with such conceptions there is general consensus that our empathic understanding and resonance with others can be elicited via two primary mechanisms: An implicit bottom-up route, and a more effortful top-down route (Batson et al., 1997; Coplan & Goldie, 2012; Decety & Jackson, 2004; Ickes, 1997; de Waal & Preston, 2017). Affective empathy is associated largely with the bottom-up route, and processes such as spontaneous facial mimicry (sometimes referred to as ‘motor resonance’) can play an important role in this component of empathy (Chartrand & van Baaren, 2009; Niedenthal, 2007; Preston & de Waal, 2001). In contrast, cognitive empathy involves more top-down, inferential processes, such as that of inhibiting one’s own perspective and taking the perspective of the other (O’Connell, Christakou, & Chakrabarti, 2015). This route requires sufficient cognitive resources to hold two representations simultaneously and to inhibit one’s default egocentric perspective in order to focus upon the other’s perspective (Chakrabarti & Baron-Cohen, 2006; Coplan & Goldie, 2012).

Emotion regulation comprises different forms, which include cognitive and inhibitory processes (Gross, 1998). One of the most widely studied forms of emotion regulation is cognitive reappraisal (henceforth reappraisal), which refers to a change in the meaning adhered to an emotion-eliciting stimulus or event in order to lessen (or increase) its emotional impact (Gross, 1998, 2001, 2002; Sheppes & Gross, 2011). Greater habitual use of reappraisal has been found to be positively related to levels of trait positive affect and metrics of social functioning (Gross & John, 2003; Nezlek & Kuppens, 2008), and negatively related to trait negative affect and propensity to develop certain clinical mood disorders (Aldao, Nolen-Hoeksema, & Schweizer, 2010; Hu et al., 2014). While paradigms to assess reappraisal often focus upon more deliberate/explicit forms of reappraisal (i.e. where participants are instructed to reappraise an emotion-eliciting stimulus in order to regulate their affective response), day-to-day emotion regulation also involves more implicit and unintentional processes (Gyurak, Gross, & Etkin, 2011; Mauss, Cook, Cheng, & Gross, 2007; Thompson, 2011). Extrinsic contextual factors can influence one’s emotional response to a stimulus in a relatively implicit manner, without any conscious goal or effort to regulate (Berkman & Lieberman, 2009; Mocaiber et al., 2011; Wang et al., 2017). In this manuscript, we use the term “explicit” to refer to instances in which reappraisal was driven by a deliberate attempt to alter one’s affective experience. We use the term “implicit” to refer to less deliberate forms of reappraisal, where one’s emotional response may be influenced by extrinsic factors without there necessarily being any conscious intent to engage in emotion regulation (Berkman & Lieberman, 2009).

The emotion state generated in the observer by perceiving another individual’s state is a function of the observer’s level of empathy. Crucially, the observer’s emotion is subject to his/her own emotion regulatory processes. The outcome of these regulatory processes, which may be activated spontaneously, often determines the resultant emotional state and response of the observer (see figure 1). Indeed, previous theoretical work has proposed a critical role for emotion regulation in empathy (Decety & Jackson, 2004; Zaki, 2014); the core assertion being that individuals with a lower capacity for emotion regulation experience higher levels of personal distress, and lower levels of other-focused sympathy when observing or imagining another individual’s negative emotional state (Eisenberg & Fabes, 1991). Consistent with this assertion, it has been shown that the relationship between trait affective empathy and self-reported prosocial behaviours is moderated by the habitual use of reappraisal (Lockwood, Seara-Cardoso, & Viding, 2014).

**Figure 1.**
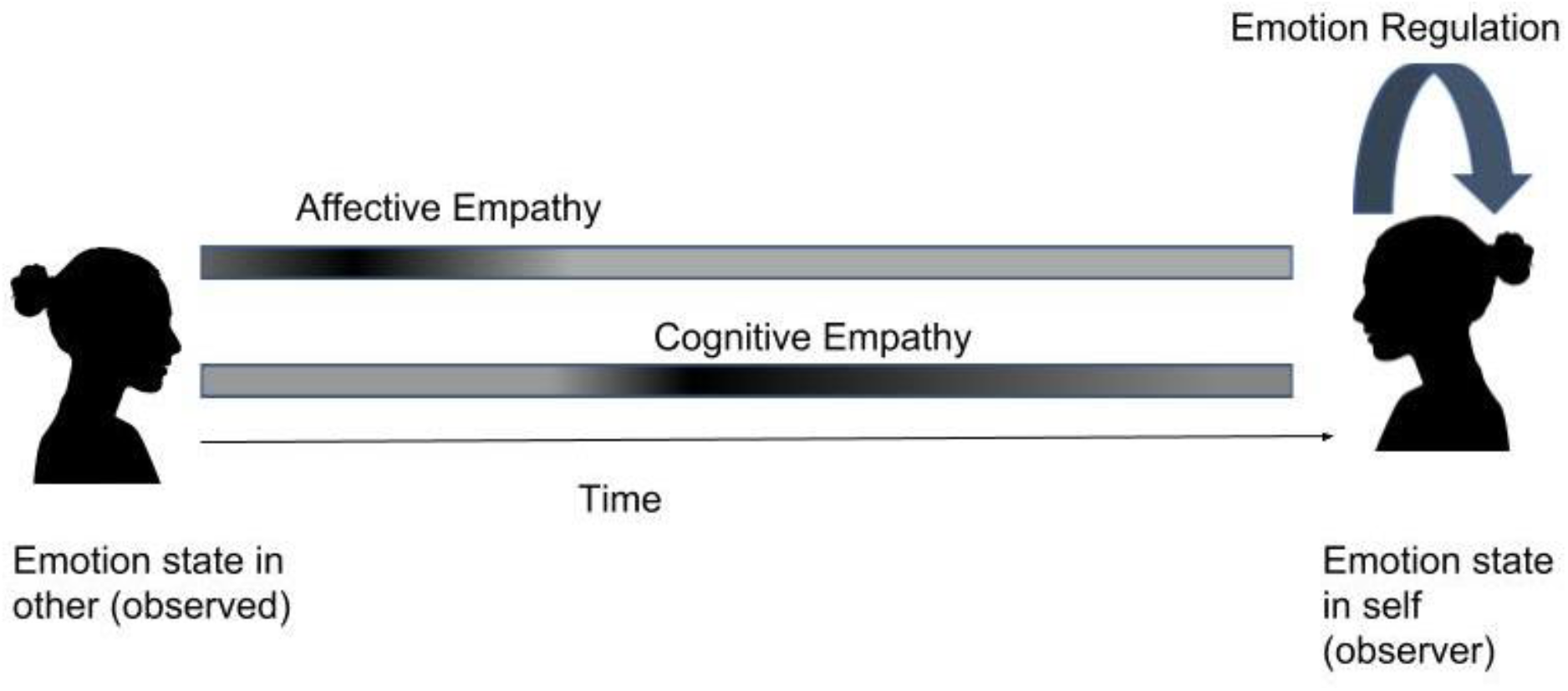
Depiction of theoretical relationship between empathy and emotion regulation. The capacity to understand and resonate with the emotional states of others is a function of the observer’s levels of cognitive and affective empathy. The colour gradient reflects the time-course of empathic processes; processes related to affective empathy occur relatively early in response to the perception of another’s emotion, with cognitive empathy processes typically coming online at a later stage. The emotion state triggered in the observer based on their comprehension and resonance with the other’s state is subject to the observer’s own emotion regulation processes.

Findings from studies in atypical populations provide additional clues on the inter-relationship of empathy and emotion regulation. Borderline personality disorder (BPD) is characterised by difficulties in regulating emotions and is also often related to atypically high levels of affective empathy (Fertuck, Lenzenweger, Clarkin, Hoermann, & Stanley, 2006; Lang et al., 2011). Individuals with autism spectrum conditions (ASC) often exhibit poor performance on measures of cognitive empathy, and also show evidence of higher levels of emotion dysregulation (Konstantareas & Stewart, 2006) and increased use of maladaptive regulation strategies (Samson, Huber, & Gross, 2012). Such findings suggest that the cognitive and affective dimensions of empathy may be differentially related to emotion regulation.

Most studies that have examined the relationship between empathy and emotion regulation have used questionnaire measures of both constructs (Lockwood et al., 2014; Eisenberg, Fabes, Guthrie, & Reiser, 2000; Okun, Shepard, & Eisenberg, 2000; Tully, Ames, Garcia, & Donohue, 2016). Given the lack of convergence between questionnaire and task-based measures often observed in studies of socio-emotional constructs (Melchers, Montag, Markett, & Reuter, 2015), we decided to explore the relationship between empathy and reappraisal using a multi-method approach combining questionnaire and task-based measures of reappraisal. The two studies reported here seek to systemically elucidate the nature of the relationship between cognitive/affective empathy and reappraisal. In study 1, we examine how measures of trait cognitive and affective empathy relate to the habitual use of reappraisal. In study 2 we examine how the same empathy measures relate to the magnitude of implicit reappraisal of negative emotional images in a behavioural task.

## Study 1

Most previous studies have focused upon the moderating effect of regulation on the association between empathy and different behavioural or clinical outcomes, such as sympathy/personal distress (Okun et al., 2000), prosocial behaviours (Lockwood et al., 2014), and depression (Tully et al., 2016). While not the key question addressed by such studies, they have provided some evidence of a direct relationship between component processes of empathy and emotion regulation. For example, Lockwood et al. (2014) found that trait cognitive, but not affective, empathy was positively correlated with the self-reported use of reappraisal. Similarly, another study found that higher levels of trait perspective taking were associated with increased use of reappraisal (Tully et al., 2016).

As reappraisal is reliant upon cognitive control processes (McCrae et al., 2010; Urry, van Reekum, Johnstone, & Davidson, 2009), it is critical that one has sufficient cognitive resources available in times of need to be able to utilise this regulation strategy effectively. This demand becomes increasingly important when considering the fact that affective experiences and exposure to emotional stimuli can have a detrimental effect on cognitive control abilities (Eysenck & Derakshan, 2011; Tottenham, Hare, & Casey, 2011). Cognitive empathy is similarly reliant upon cognitive control processes in order to inhibit one’s default self-perspective and take another’s perspective (Carlson, Mandell, & Williams, 2004; Hansen, 2011; Joseph & Tager-Flusberg, 2004; Sabbagh, Xu, Carlson, Moses, & Lee, 2006). Given the overlap in cognitive control processes, it is possible that higher levels of cognitive empathy would be associated with improved efficiency/efficacy of the cognitive processes that also support reappraisal.

While cognitive empathy may support the use of reappraisal, the opposite could in fact be true of affective empathy. Affective empathy is associated with increased reactivity to others’ emotions. Those high in affective empathy exhibit greater spontaneous facial mimicry (Lee et al., 2008; Sonnby-Borgstrom, 2002), which in turn has been shown to relate to increased self-reported resonance with the mimicked emotion (Hatfield, Cacioppo, & Rapson, 1993; Laird et al., 1994; Strayer, 1993; Wild, Erb, & Bartels, 2001). There is also evidence that affective empathy is related to increased emotional reactivity in a more general sense (Rueckert, Branch, & Doan, 2011). In contrast to cognitive empathy, the heightened arousal tendency associated with higher affective empathy may result in greater emotional interference on cognitive control, thereby reducing the individual’s capacity to engage in adaptive regulation strategies such as reappraisal. This first study tested the relationship between trait cognitive and affective empathy and self-reported habitual use of reappraisal. Based on the existing literature, we predicted that cognitive empathy would be positively related to the habitual use of reappraisal. As a corollary, we expected to find a negative relationship between affective empathy and reappraisal use.

## Method

### Participants

An a priori sample size estimation was conducted using G*Power 3.1 (Faul, Erdfelder, Lang, & Buchner, 2007). Based on a correlation coefficient of *r* = .33 between reappraisal use and trait cognitive empathy in a previous study (Lockwood et al., 2014), a minimum sample size of 67 was required to obtain power of .80. As the survey was completed online, and online data collection is typically associated with greater data loss than lab-based studies, we aimed to oversample by approximately 30% to ensure sufficient power after removal of statistical outliers and any cases with incomplete data. For reasons of convenience, data were collected from both the UK and Denmark. We aimed to collect two samples of approximately 100 participants. While no specific hypotheses were made regarding differences across these two sub-samples, we checked for consistency of patterns across samples. In total, 220 participants (161 female) were recruited from in and around the campuses of the University of Reading, UK (N = 94) and Aarhus University, Denmark (N = 126). Recruitment was via online and campus-based advertisements. All questionnaires were completed online in English, and participants were reimbursed in the form of course credits or enrolment in a lottery to win free cinema tickets. The mean age of the overall sample was 21.96 (*SD*=5.67; range=18-56). Ethical approval was obtained from the research ethics committees of the Universities of Reading and Aarhus, and informed consent was provided by all participants.

## Materials

### Empathy

Trait empathy was measured using the Questionnaire of Cognitive and Affective Empathy (QCAE; Reniers, Corcoran, Drake, Shryane, & Vollm, 2011). The QCAE is a 31-item self-report questionnaire measuring an individual’s capacity to understand and resonate with the emotions of others. It consists of five sub-components which track onto the two core facets of empathy (cognitive & affective). The cognitive empathy dimension taps into one’s propensity to take another’s perspective and accurately infer their state (e.g. “When I am upset at someone, I usually try to ‘put myself in his shoes’ for a while”, “I can sense if I am intruding, even if the other person does not tell me”). The affective empathy dimension consists of items examining the extent to which one shares in the emotions of others (e.g. “I am happy when I am with a cheerful group and sad when the others are glum”, “It affects me very much when one of my friends seems upset”). Participants rate their response to each item of the QCAE using a 4-point scale, ranging from “strongly disagree” to “strongly agree”. Cronbach’s alpha within our total sample was high for both QCAE dimensions (N=220, *α*_Cognitive Empathy_ = .90; *α*_Affective Empathy_ = .84). Cronbach’s alpha for CE and AE were similarly high within the UK and Denmark subsamples.

### Reappraisal use

Reappraisal use was measured using the Emotion Regulation Questionnaire (ERQ; Gross & John, 2003). The ERQ is a 10-item self-report questionnaire measuring the habitual propensity to use two emotion regulation strategies, namely, reappraisal and suppression. For the purposes of this study, we examined only the reappraisal dimension, which includes items such as “When I want to feel less negative emotion, I change the way I’m thinking about the situation”. Participants rate each item on a 7-point scale, ranging from “strongly disagree” to “strongly agree”. Cronbach’s alpha for the reappraisal subscale of the ERQ was high within our overall sample (N=220, *α* = .85), with similarly high alpha levels observed within each subsample.

### Data Reduction & Analyses

Normality of each measure was assessed using Kolmogorov Smirnov tests. ERQ-Reappraisal and QCAE-CE showed significant deviation from normality (ERQ-Reappraisal K-S: *D*(220) = .11, *p* <.001; QCAE-CE K-S: *D*(220) = .09, *p* <.001). The AE sub-scale of the QCAE was normally distributed (*D*(220) = .05, *p* = .20). Spearman’s rho is reported for correlations where at least one variable was non-normal. All correlations are reported as two-tailed, with a significance threshold of *p* <.05. To ensure the observed results were not overly influenced by any outlier cases, univariate and bivariate outliers were identified using a criterion of 3* interquartile range (IQR) and Cook’s D greater than 4/N, respectively. The full sample analyses are reported in the results section; results following outlier removal are reported in supplementary materials (table S1). A consistent pattern of results were observed for both analyses.

## Results

As predicted, cognitive empathy was significantly positively correlated with habitual use of reappraisal, *rho*(218) = .25, *p* <.001. In contrast, there was no relationship between affective empathy and the use of reappraisal, *rho*(218) = .06, *p* = .42 (see figure 2). These two correlations were significantly different (Steiger’s *Z* = 2.4, *p* = .02). A consistent pattern of results was observed within both sub samples: UK, CE-Reappraisal *rho*(92) = .29, *p* = .004, AE-Reappraisal *rho*(92) = .02, *p* = .88; Denmark, CE-Reappraisal *rho*(124) = .21, *p* = .02; AE-Reappraisal *rho*(124) = .09, *p* = .34.

**Figure 2.**
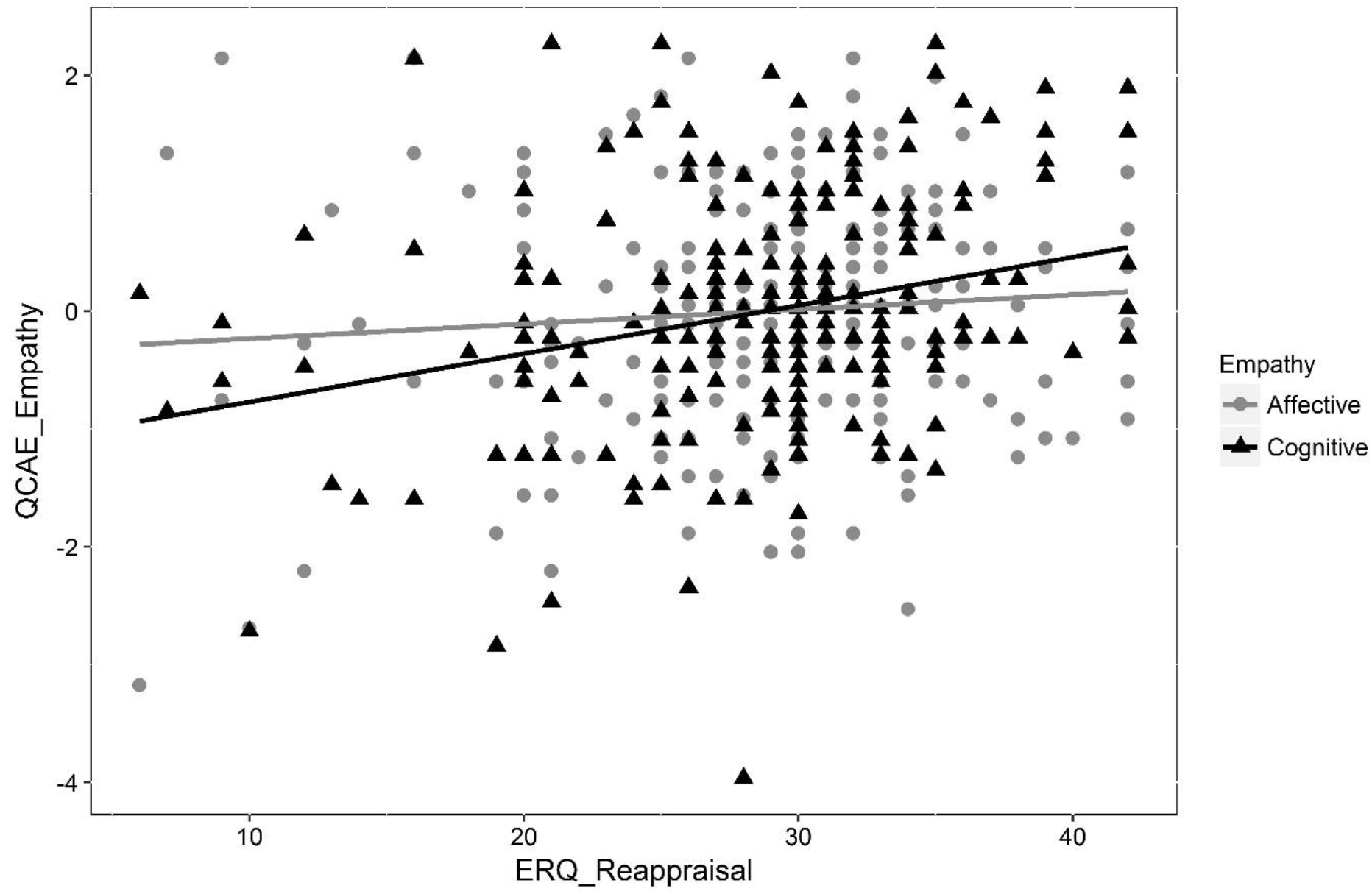
Scatterplot showing the relationship between habitual reappraisal use (ERQ) and Z-transformed cognitive/affective empathy (QCAE). Trait cognitive empathy (black) showed a positive correlation with reappraisal use. Trait affective empathy (grey) showed no relationship with reappraisal use.

These results suggest that while trait cognitive and affective empathy are positively related to one another (*rho*(218) = .28, *p* <.001), they share different relationships with the habitual use of reappraisal. Individuals with higher trait cognitive empathy report more frequent use of reappraisal in daily life. In contrast, one’s level of trait affective empathy shows no relation with the habitual use of reappraisal. Given the reliance on retrospective reporting of reappraisal use, the ERQ likely reflects participants’ tendency to use more explicit forms of reappraisal (i.e. where there is an overt intrinsic goal to alter one’s emotion, and the requirement to self-generate possible reappraisal narratives). In summary, these results suggest that cognitive, but not affective empathy, is associated with greater deliberate use of reappraisal in daily life.

## Study 2

In study 1 we observed that trait cognitive and affective empathy show different relationships with the habitual use of reappraisal. To investigate this result further, in this second study we examined the relationship between trait empathy and a task-based measure of reappraisal. The majority of task-based measures of reappraisal have utilised what is often referred to as explicit (or instructed) reappraisal paradigms. In such tasks, participants are specifically instructed to use reappraisal to modify their emotional experience (e.g. Ochsner et al., 2004). This approach and its variants have demonstrated a reliable influence of explicit reappraisal on self-reported emotional experience, psychophysiological responses (Jackson, Malmstadt, Larson, & Davidson, 2000; Ray, McCrae, Ochsner, & Gross, 2010), and neural activation in regions associated with emotional processing and self-regulation (for reviews see Kalisch, 2009; Ochsner & Gross, 2008).

In explicit reappraisal tasks, participants are typically required to self-generate alternative appraisals of the emotional stimuli. Given the inherent cognitive demands associated with such processes (McCrae et al., 2010; Urry et al., 2009), some have questioned the extent to which the observed differences between conditions in these tasks may reflect variability in the cognitive demands inherent in these different instructions, rather than being directly attributable to the change in emotional appraisal itself (Foti & Hajcak, 2008). Other studies have adopted paradigms that aim to assess more implicitly evoked forms of reappraisal. Such tasks typically involve pairing emotional stimuli with brief descriptions (or ‘frames’) that provide a context for interpreting what is happening (e.g. Foti & Hajcak, 2008; Wang et al., 2017). In a seminal early study examining the effects of contextual framing on emotional responses, Lazarus and Alfert (1964) demonstrated how self-report and physiological indices of stress can be modulated by auditory descriptions that influence the interpretation of stress-inducing video clips. More recent studies provide further evidence of the modulatory effect of contextual framing on emotional responding (Dennis & Hajcak, 2009; Foti & Hajcak, 2008; Kim et al., 2004; MacNamara, Foti, & Hakcak, 2009; Mocaiber et al., 2011; Wang et al., 2017; Wu, Winkler, Andreatta, Hajcak, & Pauli, 2012).

In contrast to explicit reappraisal paradigms, participants are not given any explicit instruction to regulate their emotions in framing tasks and are required to simply attend to the images in all conditions. Thus, framing tasks are distinct from explicit reappraisal tasks in that their influence on emotional experience is driven by extrinsic factors, and they are thought to reflect reappraisal of a more implicit and unintentional nature (Mocaiber et al., 2011; Wang et al., 2017; Foti & Hajcak, 2008). Framing tasks control for the potential confounding effects of cognitive demand that may be inherent in explicit reappraisal tasks, and by defining an appraisal narrative they also control for variability in the specific nature of the reappraisal (MacNamara et al., 2009).

In this study we examine how variability in empathy relates to the implicit reappraisal of negative images. Implicit reappraisal was operationalised as the difference in ratings of self-reported negative experience between negative images paired with descriptive framing sentences (which simply described the image content, providing a baseline measure of participants’ emotional reaction to the negative images) and negative images paired with neutralising framing sentences (aimed to dampen the unpleasantness of the images). Given the findings from study 1, we predicted that implicit reappraisal would be positively related to trait cognitive empathy. Based upon the available literature and the findings of study 1, there was no reason to predict any relationship between affective empathy and implicit reappraisal.

## Method

### Participants

Based on a correlation coefficient of.25 reported in study 1 and a previous study reporting a coefficient of.33 (Lockwood et al., 2014), to detect a moderate correlation of.3 between trait cognitive empathy and reappraisal with power of.80, an a priori sample size estimation suggested a minimum sample of 82 was required. A sample of 92 participants (73 females) was recruited from the undergraduate population at the University of Reading to take part in a one-hour study on “mood and cognitive performance”. All participants had normal or corrected to normal vision, and the mean age was 20.24 years (*SD* = 2.21; range = 18-35). Participants were recruited via the university online SONA system, and were awarded course credit for their participation.

## Materials

### Empathy

As in study 1, we used the QCAE (Reniers et al., 2011) to measure trait empathy. Within this sample, both sub-scales of the QCAE demonstrated high internal consistency (*α*_Cognitive Empathy_ = .87; *α*_Affective Empathy_ = .84).

### Reappraisal use

The ERQ (Gross & John, 2003) was used to measure self-reported use of reappraisal. Cronbach’s alpha for the Reappraisal sub-scale within this sample was acceptable (*α* = .70).

### Affective images

Forty images were selected from the International Affective Picture System (IAPS) database (Lang, Bradley, & Cuthbert, 2005). There were 20 negative and 20 neutral images. All images were of a social nature, which we defined broadly as images involving a sentient target (or targets), for which one could infer an emotional/cognitive state, and/or experience an emotional reaction in response to observing the target’s situation. Within this broader social category, we used a range of image types from the IAPS, including: Mutilation/injury, assault/attack, soldier, firefighter, car accident, scared/sick child, drugs, burn victim, battered female (negative); neu-man/woman/child, office, bakers, factory worker, harvest (neutral).

Based on the normative ratings provided in the IAPS manual (Lang et al., 2005), independent samples t-tests demonstrated that the negative images had a significantly lower mean valence rating (*M*_*valence*_ = 2.35, SD = 0.41), and significantly higher mean arousal rating (*M*_*arousal*_ = 5.67, SD = 0.91), compared to the neutral images (*M*_*valence*_ = 5.25, *SD*_*valence*_ = 0.56; *M*_*arousal*_ = 3.56, *SD*_*arousal*_ = 0.49), (valence, *t*(38) = 18.58, *p* <.001; arousal, *t*(38) = 9.14, *p* <.001).

### Reappraisal framing sentences

Forty framing sentences were used in the study (20 neutralising, 20 descriptive). Where possible, sentences were taken from prior studies on reappraisal framing. Seven sentences were taken from Foti and Hajcak (2008); two of which were edited slightly to make the context less ambiguous. Thirty-two sentences were taken from an unpublished study conducted by collaborators, and one new sentence was created to fit our image set (details of all negative and neutral IAPS images, with accompanying framing sentences are available in supplementary material, tables S2 & S3 respectively). An independent samples t-test confirmed that the different sentence types did not differ significantly in terms of word count (*M*_*descripti8ve*_ = 8.6, *SD*_*descriptive*_ = 1.79; *M*_*neutralising*_ = 8.7, *SD*_*neutralising*_ = 1.84), *t*(38) = .261, *p* = .80.

All participants saw the same images in block 1 (freeview condition). Half of the neutral images from block 1 were shown again in block 2 with an accompanying descriptive framing sentence. The ten neutral images presented in block 2 were counterbalanced across participants. Based on the ratings provided in the IAPS manual (Lang et al., 2005), an independent samples t-test demonstrated that the different neutral image sets shown in block 2 did not differ significantly in valence, *t*(18) = .94, *p* = .36, or arousal, *t*(18) = .67, *p* = .51.

For block 2 of the task, two sets of 10 images were created from the 20 negative images shown in block 1. We tried to match both negative image sets based on factors such as the image category (e.g. injury, drugs etc.), complexity, and the age, race, and number of depicted individuals. Based on the normative ratings (Lang et al., 2005), independent samples t-tests showed that these two negative image sets did not differ significantly in terms of valence (*M*_set1_ = 2.33, *SD*_*set1*_ = 0.45; *M*_*set2*_ = 2.37, *SD*_*set2*_ = 0.39) or arousal (*M*_set1_ = 5.65, *SD*_*set1*_ = 0.62; *M*_*set2*_ = 5.70, *SD*_*set2*_ = 1.16); valence, *t*(18) = .20, *p* = .85; arousal, *t*(18) = .12, *p* = .91. In block 2, half of the negative images were paired with descriptive sentences, and half with neutralising sentences; image-sentence pairings were counterbalanced across participants.

### Procedure

Participants read the information sheet and provided written consent to participate. All testing was completed in isolation, in a distraction-free environment. Participants were sat at an approximate viewing distance of 60cm from the monitor. The task was run on a PC at a resolution of 600 × 800, with a monitor refresh rate of 60Hz. The implicit reappraisal framing task was completed as part of a larger battery, which took approximately one hour in total. The task consisted of two blocks, with 70 image trials in total (Block 1: 20 negative, 20 neutral images; Block 2: 20 negative, 10 neutral images), and lasted on average approximately 20 minutes. While our focus was upon the ratings for the negative images under different framing conditions, we included neutral images as a means of slowing habituation to negative images.

Participants were informed that they would view a series of images and were asked to report after each presentation “how unpleasant/pleasant they felt in response to the image”. Ratings were made using the keyboard and a 1-9 bipolar scale was used, where: 1 = extremely unpleasant, 5 = neutral, 9 = extremely pleasant. While no positive images were used in the task, a bipolar valence scale covering the range from unpleasant to pleasant was used to maintain consistency with ratings scales used in previous studies (Foti & Hajcak, 2008; MacNamara et al., 2009) and to account for the possibility that the neutralising reappraisal frames could result in the images being appraised as more positive/pleasant than neutral. Participants were asked to provide honest/accurate ratings based on their “initial reaction” upon viewing each image. The images in each block were presented in a randomised order. Block 1 trials followed the same sequence as those in block 2 (depicted in figure 3), except there was no framing sentence event between the initial fixation and image presentation.

**Figure 3.**
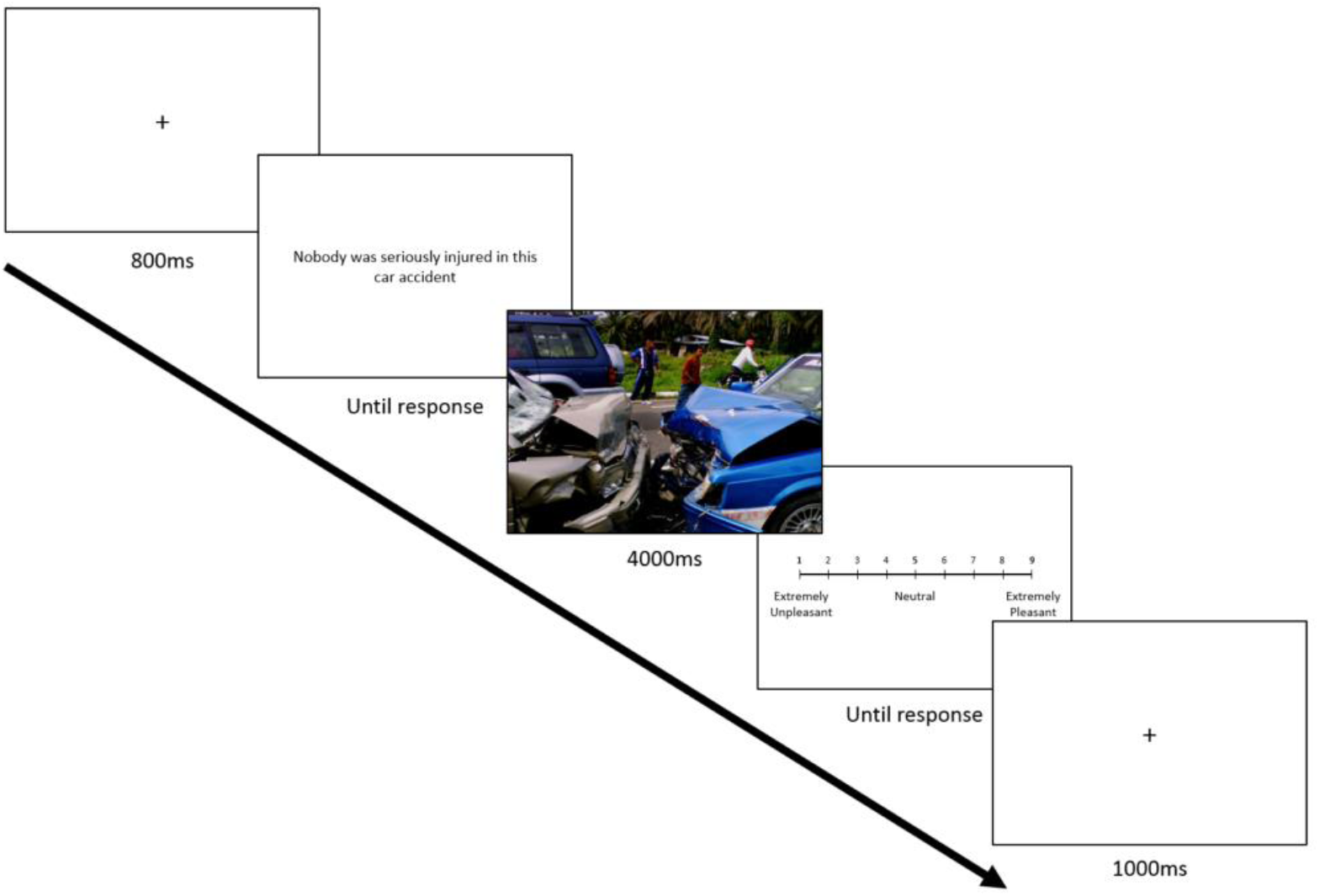
Schematic of trial events during block 2 of the implicit reappraisal task. Each trial consisted of the following events: 1) white screen with black central fixation cross (800ms); 2) neutralising or descriptive framing sentence, which remained on screen until response was made); 3) IAPS image (neg or neu; 4000ms); 4) white screen with centrally presented rating scale (1-9, remained on screen until response was made); 5) white screen with black central fixation cross (1000ms). The image shown above was selected from the OASIS images set (Kurdi, Lozano, & Banji, 2017) as a representative example and was not one of those used in the task.

Prior to commencement of block 2, participants were given the opportunity to take a short break. They were then presented with a new instruction screen which informed them that in this next block each image would be “preceded by a sentence which provides a context for what is happening in the image”. At no point prior to or during the testing session was there any reference to emotion regulation. Block 2 consisted of the same 20 negative images from block 1, with half preceded by a descriptive framing sentence and the other half preceded by a neutralising framing sentence. The 10 neutral images presented in block 2 were all preceded by a descriptive framing sentence. The term reappraisal implies a process of altering one’s interpretation of a situation after an initial appraisal has been formed. While the framing sentences preceded the images in block 2, participants had already viewed and formed an appraisal of the images during the freeview condition (block 1). As any notable change in the intensity of negative experience elicited by the images would rely upon participants altering their original appraisals, this task is best defined as a measure of implicit reappraisal, rather than what is sometimes referred to as ‘pre-appraisal’ (Matarazzo, Baldassarre, Nigro, & Abbamonte, 2014).

### Data reduction & analyses

Two participants were removed prior to analysis for failing to correctly follow task instructions, leaving a final sample of N = 90. Our key metric of implicit reappraisal was the extent to which participants reported a decreased negative emotional response to negative images paired with neutralising framing sentences (NegNEU condition), relative to negative images paired with descriptive framing sentences (NegDES condition). This metric was termed ‘implicit reappraisal’ and was calculated for each participant by subtracting their mean NegDES rating from their mean NegNEU rating. Thus, higher implicit reappraisal scores reflect a greater reduction in unpleasant experience as a result of the neutralising framing sentences. This metric provides a measure of the extent to which the extrinsic framing sentence type influenced participants’ emotional reaction to the negative images, while controlling for the intrinsic valence of the stimulus.

We conducted separate paired samples t-tests to explore the effects of image (negative, neutral) and sentence frame (neutralising, descriptive) on self-reported valence ratings. Individual differences were examined by correlations exploring the relationship between the implicit reappraisal task metric and trait empathy measures (QCAE-CE, QCAE-AE). Normality of each variable was assessed using Kolmogorov-Smirnov tests. QCAE-CE and the task measure of implicit reappraisal were normally distributed (*D*(90) = .08, *p* = .20, & *D*(90) = .09, *p* = .07, respectively). QCAE-AE showed significant deviation from normality (*D*(90) = .09, *p* = .049). All correlations are reported as two-tailed, with a significance threshold of *p* <.05. Spearman’s Rho is reported for correlations where any of the variables deviated from normality. To ensure the observed results were not overly influenced by any outlier cases, univariate and bivariate outliers were identified using a criterion of 3*IQR and Cook’s D greater than 4/N, respectively. The full sample analyses are reported in the results section; results following outlier removal are reported in supplementary materials (table S4). A consistent pattern of results were observed for both analyses.

## Results

### Questionnaire correlations

We repeated the trait correlation analyses from study 1, to test whether we could replicate the findings in a different sample. These results showed a consistent pattern to study 1: The positive correlation between cognitive empathy and reappraisal use was just on the threshold of significance, *r*(88) = .20, *p* = .057; affective empathy showed no relationship with reappraisal use, *rho*(88) = −.05, *p = .*64. These two correlations were significantly different to one another (Steiger’s *Z* = 2.1, *p* = .04).

### Implicit reappraisal task

Block 1 valence ratings for negative images (*M* = 2.49, *SD* = .77) were significantly more negative than for neutral images (*M* = 5.58, *SD* = .60), *t*(89) = −27.94, *p* <.001. Valence ratings in the NegNEU condition (*M* = 4.38, *SD* = .97) were significantly less negative than in the baseline NegDES condition (*M* = 2.92, *SD* = .77), *t*(89) = 15.17, *p* <.001. This suggests that the context-framing manipulation had the expected effect on participant’s self-reported emotional experience. See figure 4 for mean valence ratings across all conditions.

**Figure 4.**
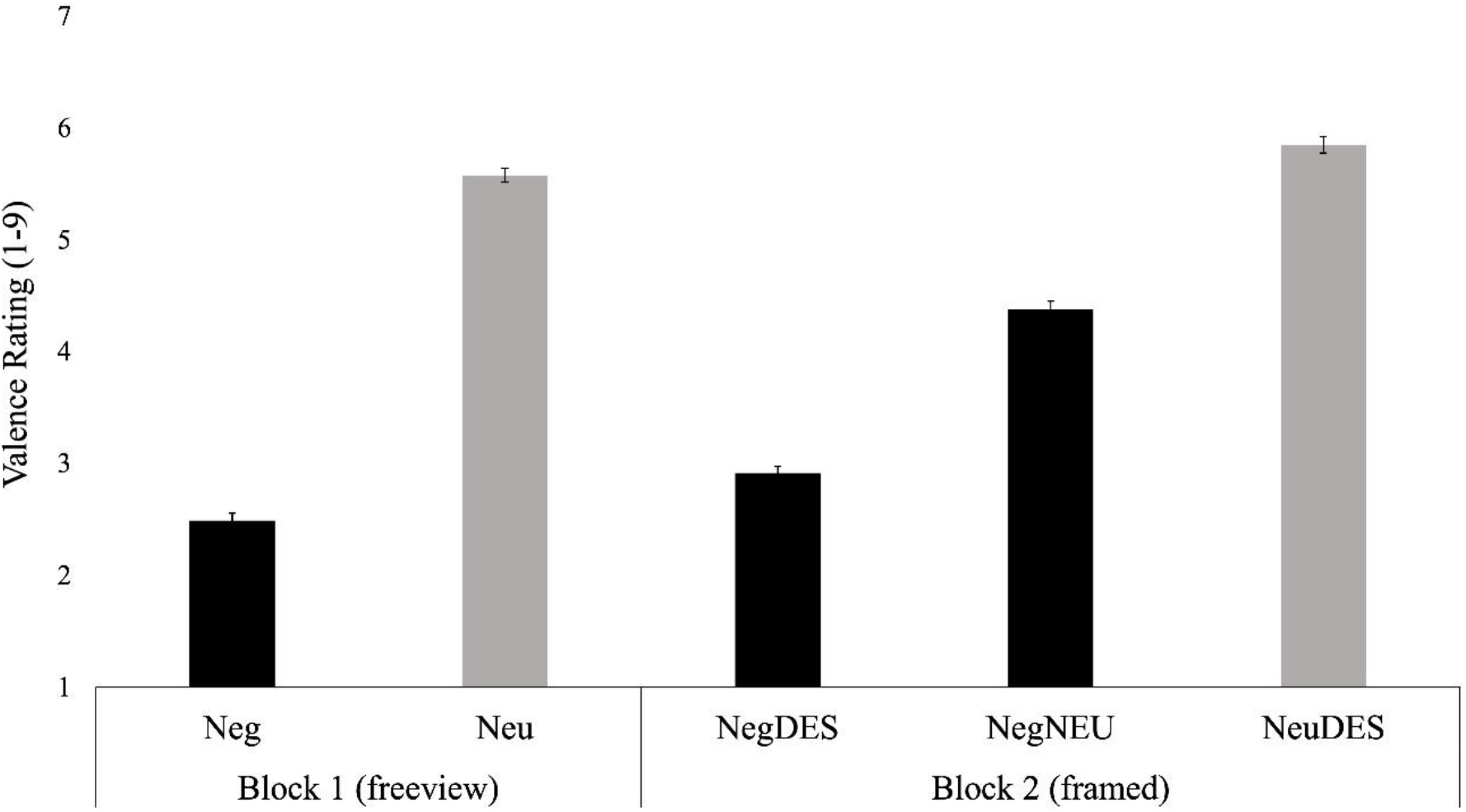
Mean valence ratings across all conditions in the implicit reappraisal task (1 = extremely unpleasant, 5 = neutral, 9 = extremely pleasant). In block 1 (freeview condition), negative images (Neg) were associated with greater self-reported negative experience relative to neutral images (Neu). In block 2 (framed condition), negative images paired with a neutralising framing sentence (NegNEU) showed reduced self-reported negative (more neutral) experience relative to the baseline condition in which negative images were paired with a descriptive framing sentence (NegDes). Error bars depict ±1 within-subjects SEM.

### Effects of empathy on implicit reappraisal

Trait cognitive empathy showed no relationship with the implicit reappraisal metric (NegNEU minus NegDES), *r*(88) = −.06, *p* = .60. However, there was a positive correlation between affective empathy and implicit reappraisal, *rho*(88) = .33, *p* = .002; higher levels of affective empathy were associated with reduced negative/more neutral ratings of negative images after providing a neutralising framing context (see figure 5). These two correlations were significantly different (Steiger’s *Z* = −3.22, *p* = .001). As in study 1, while trait cognitive and affective empathy were positively correlated with one another (*rho*(88) = .36, *p* <.001), they showed different relationships with our reappraisal measure. The results of this study suggest that affective, but not cognitive, empathy is related to the capacity to implicitly reduce negative emotional experience in response to the neutralising context frames.

**Figure 5.**
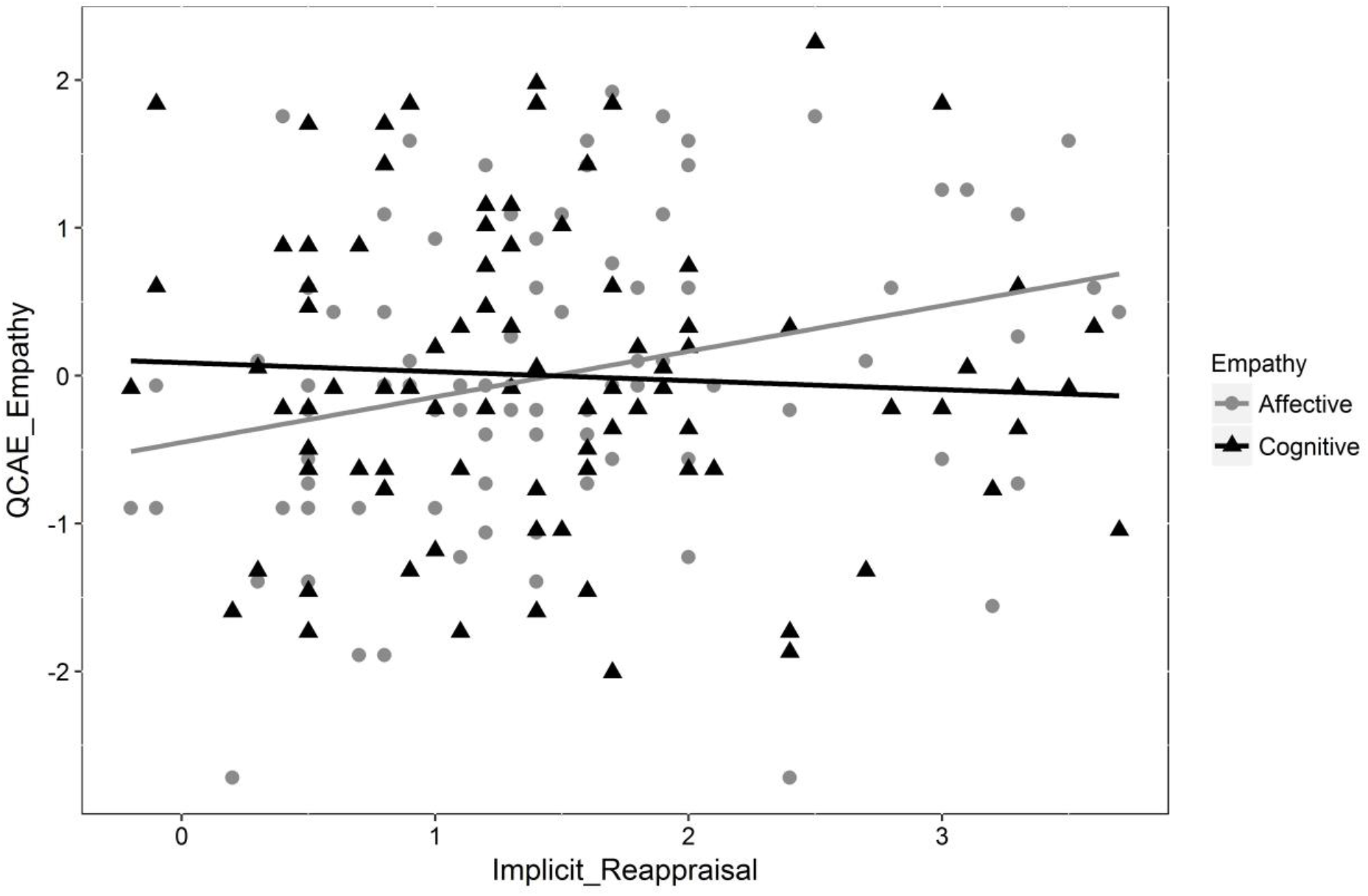
Scatterplot showing the relationship between the implicit reappraisal task metric (NegDES - NegNEU condition valence ratings) and Z-transformed trait cognitive/affective empathy (QCAE). Affective empathy (grey) was positively correlated with implicit reappraisal (i.e. enhanced reduction in self-reported negative experience for negative images paired with a neutralising framing sentence, relative to negative images paired with a descriptive framing sentence). Trait cognitive empathy (black) showed no relationship with implicit reappraisal.

## Discussion

We report two studies in which questionnaire and task measures were used to examine the relationship between trait cognitive/affective empathy and reappraisal. Consistent with prior work asserting a distinction between the two facets of empathy (Coplan & Goldie, 2012; Singer & Lamm, 2009), in both studies we observed different relationships between cognitive/affective empathy and reappraisal, depending on the way reappraisal was measured. Across both studies, cognitive empathy was positively related to the habitual use of reappraisal, and there was no evidence of a relationship between affective empathy and reappraisal use. This finding replicates prior work, which has reported the same relationships between these two dimensions of empathy and reappraisal use (Lockwood et al., 2014). During the preparation of this manuscript, a study by Powell (2018) was published, which reported the same relationships between reappraisal use and trait cognitive and affective empathy in a large UK sample. The findings of our study provide further support for this recent work by replicating the findings twice, including in a sample from two countries, and demonstrating that the relationships between cognitive/affective empathy and reappraisal use are significantly different.

In study 2, the finding of a significant reduction in self-reported unpleasant experience for negative stimuli paired with neutralising framing sentences is consistent with prior work demonstrating an influence of context framing on emotional experience (Dennis & Hajcak, 2009; Foti & Hajcak, 2008; Kim et al., 2004; MacNamara et al., 2009; Mocaiber et al., 2011; Wang et al., 2017; Wu et al., 2012). In contrast to the results of study 1, it was observed that affective, but not cognitive, empathy was positively related to implicit reappraisal (reflected by the extent to which participants experienced a reduction in negative emotions for negative images preceded by neutralising frames, relative to descriptive frames). Taken together, these findings suggest that while higher cognitive empathy is associated with increased reappraisal use in daily life, it does not provide any significant advantage in implicit reappraisal driven by extrinsic contextual information. Similarly, while affective empathy is not related to one’s propensity to use reappraisal in daily life, it may support the ability to alter initial emotional appraisals on a more implicit level. While the correlational design of our studies precludes the ability to make cause and effect judgments, below we discuss some possible interpretations of these results in relation to prior findings and theory.

The positive relationship between trait cognitive empathy and the habitual use of reappraisal suggests that higher levels of cognitive empathy are associated with an increased propensity to use reappraisal in daily life. The ERQ measure of reappraisal is based on retrospective self-report, and therefore likely reflects the use of more deliberate and intrinsically driven forms of reappraisal. Explicit reappraisal and cognitive empathy rely upon similar cognitive control processes, such as attentional switching, working memory, and inhibitory control, and are both related to activation in partially overlapping regions of the PFC (Kalisch, 2009; McCrae et al., 2010; Sabbagh et al., 2006; Urry et al., 2009). Enhanced cognitive control abilities associated with higher levels of cognitive empathy might therefore facilitate one’s ability, and thus propensity, to utilise reappraisal in a deliberate and conscious manner to regulate emotions.

While cognitive empathy was associated with the habitual use of reappraisal, it showed no relationship with downregulation of negative emotion in a context-framing task. The framing task provided a measure of more implicit reappraisal, specifically, the downregulation of negative affect resulting from extrinsic contextual information, without a specific instruction or conscious intent to alter one’s emotional state. Such implicit reappraisal processes are less cognitively demanding than reappraisal use in many real-life situations, as there is no need for explicit goal switching and/or self-generation of reappraisal narratives (Wang et al., 2017). Thus, it might be that high cognitive empathy supports only the more explicit, cognitively demanding forms of reappraisal and offers little advantage in terms of more implicit reappraisal processes. Such an interpretation could explain why higher trait cognitive empathy was related to more frequent use of reappraisal in daily life but was not related to downregulation of negative emotions in our implicit reappraisal task.

While affective empathy was not related to self-reported frequency of reappraisal use, individuals with higher levels of affective empathy showed an increased magnitude of implicit reappraisal in the framing task. This suggests that affective empathy may support the ability to implicitly alter one’s emotional appraisals based on new contextual information but is unrelated to one’s propensity to explicitly use reappraisal in daily life. While there is overlap in the brain regions associated with explicit and implicit reappraisal, explicit reappraisal represents a more cognitively demanding task (due in part to the recruitment of overt goal switching and semantic processes to construct potential reappraisal narratives) (Ochsner & Gross, 2005; Wang et al., 2017). Reappraisal frames are thought to exert their influence over emotional responses in a relatively implicit manner (Foti & Hajcak, 2008; Wang et al., 2017). Affective empathy is similarly thought to involve largely implicit processing of emotional information (Chartrand & van Baaren, 2009; Niedenthal, 2007; Preston & de Waal, 2001). Hence, the association between implicit reappraisal and affective empathy is not entirely surprising given that both abilities relate to implicit emotional processes.

It is important to consider a caveat of the current set of studies that is shared with many studies on emotion regulation. While we adopted both questionnaire and task-based measures of reappraisal, all of our key metrics were based on self-report. Socially desirable responding and/or demand characteristics could have played some role in the observed effects. A number of steps were taken to reduce the potential for such confounds: Participants were recruited for a study on “mood and cognitive performance”, and there was no mention of emotion regulation during the lab testing session. Furthermore, the experimenter was not present in the room during task completion, and participants were explicitly informed that their data would be analysed anonymously and were instructed to respond accurately and honestly.

The findings of these studies represent a further step towards understanding the nature of the relationship between cognitive/affective empathic processes and reappraisal, and may have broader implications, for example, in helping to understand the nature of the shared empathy and emotion regulation deficits observed in psychopathological groups, such as in individuals with ASC. An important avenue for further research is to examine the relationship between empathy and emotion regulation abilities using more objective measures of emotional reactivity/regulation, such as facial EMG, EEG and functional MRI based approaches, as well as examining how empathy relates to other regulation strategies such as expressive suppression or distraction (Gross, 2015). Further, given the lack of convergence between trait and task measures of empathy observed in other studies (Melchers et al., 2015), it would be valuable to test the relationships observed in this study using a broader range of measures of cognitive and affective empathy, and by studying both explicit and implicit emotion regulation using tasks and questionnaires (Gyurak et al., 2011). Finally, as our sample consisted primarily of typically developing university students, it would also be worthwhile to study this relationship in additional samples, such as children, older adults, and atypical populations.

## Acknowledgements

This research is supported by a Medical Research Council (MRC) doctoral studentship (MR/J003980/1) awarded to Nicholas M. Thompson.

## Contributions

NMT contributed to study design, data collection/analysis, and manuscript preparation. CVM and BC contributed to study design and manuscript preparation.

## Data availability statement

The data from these studies are available upon request to the corresponding author.

## References

Aldao, A., Nolen-Hoeksema, S., Schweizer, S. (2010). Emotion-regulation strategies across psychopathology: A meta-analytic review. Clinical Psychology Review, 30: 217–237.

Baron-Cohen, S., & Wheelwright, S. (2004). The empathy quotient: an investigation of adults with Asperger syndrome or high functioning autism, and normal sex differences. Journal of Autism & Developmental Disorders, 34: 163–175.

Batson, D. C., Sager, K., Garst, E., Kang, M., Rubchinsky, K., & Dawson, K. (1997). Is empathy-induced helping due to self-other merging? Journal of Personality & Social Psychology, 73: 495-509.

Berkman, E. T., & Lieberman, M. D. (2009). Using neuroscience to broaden emotion regulation: Theoretical and methodological considerations. Social & Personality Psychology Compass, 3: 475–493.

Carlson, S. M., Mandell, D. J., & Williams, L. (2004). Executive function and theory of mind: stability and prediction from ages 2 to 3. Developmental Psychology, 40: 1105–22.

Chakrabarti, B., & Baron-Cohen, S. (2006). Empathizing: Neurocognitive developmental mechanisms and individual differences. Progress in Brain Research, 156: 403–417.

Chartrand, T. L., & van Baaren, R. (2009). Human mimicry. Advances in Experimental Social Psychology, 41: 219–274.

Coplan, A., & Goldie, P. (2012). Two Routes to Empathy. Oxford Press.

Decety, J., & Jackson, P. L. (2004). The functional architecture of human empathy. Behavioral & Cognitive Neuroscience Reviews, 3: 71–100.

Decety, J., & Lamm, C. (2006). Human empathy through the lens of social neuroscience. The Scientific World, 6: 1146–1163.

Dennis, T. A., & Hajcak, G. (2009). The late positive potential: a neurophysiological marker for emotion regulation in children. Journal of Child Psychology & Psychiatry, 50: 1373–1383.

de Waal, F. B. M., & Preston, S. D. (2017). Mammalian empathy: behavioural manifestations and neural basis. Nature Reviews Neuroscience, 18: 498–509.

Eisenberg, N., & Fabes, R. A. (1991). Prosocial behavior and empathy: a multimethod, developmental perspective. In M Clark (Ed.), Review of Personality and Social Psychology, 12: 34–61. CA: Sage.

Eisenberg, N., Fabes, R. A., Guthrie, I. K., & Reiser, M. (2000). Dispositional emotionality and regulation: their role in predicting quality of social functioning. Journal of Personality & Social Psychology, 78: 136–57.

Eysenck, M. W., & Derakshan, N. (2011). New perspectives in attentional control theory. Personality & Individual Differences, 50: 955–960.

Faul, F., Erdfelder, E., Lang, A. G., & Buchner, A. (2007). G*Power 3: a flexible statistical power analysis program for the social, behavioral, and biomedical sciences. Behaviour Research Methods, 39:175–91.

Fertuck, E. A., Lenzenweger, M. F., Clarkin, J. F., Hoermann, S., & Stanley, B. (2006). Executive neurocognition, memory systems, and borderline personality disorder. Clinical Psychology Review, 26: 346–375.

Fodor, J. A. (1987). Psychosemantics: The Problem of Meaning in the Philosophy of Mind. MA: MIT Press.

Foti, D., Hajcak, G., 2008. Deconstructing reappraisal: descriptions preceding arousing pictures modulate the subsequent neural response. Journal of Cognitive Neuroscience, 20: 977–988.

Gross, J.J. (1998). The emerging field of emotion regulation: an integrative review. Review of General Psychology, 2: 271–299.

Gross, J.J. (2001). Emotion regulation in adulthood: timing is everything. Current Directions in Psychological Science 10: 214–219.

Gross, J. J. (2002). Emotion regulation: Affective, cognitive, and social consequences. Psychophysiology, 39: 281–291.

Gross, J. J. (2015). Emotion Regulation: Current Status and Future Prospects. Psychological Inquiry: An International Journal for the Advancement of Psychological Theory, 26: 1–26.

Gross, J. J., & John, O. P. (2003). Individual differences in two emotion regulation processes: implications for affect, relationships, and well-being. Journal of Personality & Social Psychology, 85: 348–362.

Gyurak, A., Gross, J.J., Etkin, A. (2011). Explicit and implicit emotion regulation: a dual-process framework. Cognition & Emotion, 25, 400–412.

Hansen, S. (2011). Inhibitory control and empathy-related personality traits: Sex-linked associations. Brain and Cognition, 76: 364–368.

Hatfield, E., Cacioppo, J.T., & Rapson, R.L. (1993) Emotional contagion. Current Directions in Psychological Science, 2: 96–99.

Hu, T., Zhang, D., Wang, J., Mistry, R., Ran, G., & Wang, X. (2014). Relation between emotion regulation and mental health: A meta-analysis review. Psychological Reports, 114: 341–62.

Ickes, W. (1997) Empathic Accuracy. New York: The Guilford Press.

Jackson, D. C., Malmstadt, J. R., Larson, C. L., & Davidson, R. J. (2000). Suppression and enhancement of emotional responses to unpleasant pictures. Psychophysiology, 37: 515–22.

Joseph, R. M., & Tager-Flusberg, H. (2004). The relationship of theory of mind and executive functions to symptom type and severity in children with autism. Development & Psychopathology, 16: 137–55.

Kalisch, R. (2009). The functional neuroanatomy of reappraisal: Time matters. Neuroscience & Biobehavioral Reviews, 33: 1215–1226.

Kim, H., Somerville, L. H., Johnstone, T., Polis, S., Alexander, A. L., Shin, L. M., & Whalen PJ. (2004). Contextual modulation of amygdala responsivity to surprised faces. Journal of Cognitive Neuroscience., 16: 1730–45.

Konstantareas, M. M., & Stewart, K. (2006). Affect regulation and temperament in children with Autism Spectrum Disorder. Journal of Autism & Developmental Disorders, 36: 143–54.

Kurdi, B., Lozano, S., & Banji, M. R. (2017). Introducing the Open Affective Standardized Image Set (OASIS). Behaviour Research Methods, 49: 457–470.

Laird, J. D., Alibozak, T., Davainis, D., Deignan, K., Fontanella, K., Hong, J., Levy, B., & Pacheco, C. (1994). Individual differences in the effects of spontaneous mimicry on emotional contagion. Motivation & Emotion, 18: 231–247.

Lang, P.J., Bradley, M.M., & Cuthbert, B.N. (2005). *International affective picture system* (IAPS): Affective ratings of pictures and instruction manual. Technical Report A-6. University of Florida, Gainesville, FL.

Lang, S., Stopsack, M., Kotchoubey, B., Frick, C., Grabe, H. J., Spitzer, C., & Barnow, S. (2011). Cortical inhibition in alexithymic patients with borderline personality disorder. Biological Psychology, 88: 227–232.

Lazarus, R.S., Alfert, E. (1964). Short-circuiting of threat by experimentally altering cognitive appraisal. Journal of Abnormal & Social Psychology, 69: 195–205.

Lee, B. T., Seok, J. H., Lee, B. C., et al. (2008). Neural correlates of affective processing in response to sad and angry facial stimuli in patients with major depressive disorder. Progress in Neuro-Psychopharmacology & Biological Psychiatry, 32: 778–85.

Lockwood, P. L., Seara-Cardoso, A., & Viding, E. (2014). Emotion regulation moderates the association between empathy and prosocial behavior. Plos One, 9.

MacNamara, A., Foti, D., Hajcak, G. (2009). Tell me about it: neural activity elicited by emotional pictures and preceding descriptions. Emotion, 9: 531–543.

Matarazzo, O., Baldassarre, I., Nigro, G., Abbamonte, L. (2014). Helpful contextual information before or after negative events: Effects on appraisal and emotional reaction. Cognitive Computation 6: 640–651.

Mauss, I. B., Cook, C. L., Cheng, J. Y. J., & Gross, J. J. (2007). Individual differences in cognitive reappraisal: Experiential and physiological responses to an anger provocation. International Journal of Psychophysiology, 66: 116–24.

McCrae, K., Hughes, B., Chopra, S., John, G., Gross, J.J., & Ochsner, K. N. (2010). The Neural Bases of Distraction and Reappraisal. Journal of Cognitive Neuroscience, 22: 248–262.

Melchers, M., Montag, C., Markett, S., & Reuter, M. (2015). Assessment of empathy via self-report and behavioural paradigms: data on convergent and discriminant validity. Cognitive Neuropsychiatry, 20: 157–71.

Mocaiber, I., Sanchez, T. A., Pereira, M. G., Erthal, F. S., Joffily, M., Araijjo, D. B., Volchan, E., De Oliveira, L. (2011). Antecedent descriptions change brain reactivity to emotional stimuli: A functional magnetic resonance imaging study of an extrinsic and incidental reappraisal strategy. Neuroscience, 193: 241–248.

Nezlek, J. B., & Kuppens, P. (2008). Regulating positive and negative emotions in daily life. Journal of Personality, 76: 561–80.

Niedenthal, P. M. (2007). Embodying emotion. Science, 18: 1002–5.

Ochsner, K. N. and Gross, J. J. (2005). The cognitive control of emotion. Trends in Cognitive Sciences, 9: 242–249.

Ochsner, K. N., Gross, J. J. (2008). Cognitive emotion regulation: insights from social cognitive and affective neuroscience. Current Directions in Psychological Science, 17: 153–158.

Ochsner, K. N., Ray, R. D., Cooper, J. C., Robertson, E. R., Chopra, S., Gabrieli, J. D., & Gross, J. J. (2004). For better or for worse: neural systems supporting the cognitive down- and up-regulation of negative emotion. Neuroimage, 23: 483–99.

O’Connell, G., Christakou, A. and Chakrabarti, B. (2015). The role of simulation in intertemporal choices. Frontiers in Neuroscience, 9: 1662–453.

Okun, M. A., Shepard, S. A., & Eisenberg, N. (2000). The relations of emotionality and regulation to dispositional empathy-related responding among volunteers-in-training. Personality & Individual Differences, 28: 367–382.

Powell, P. A. (2018). Individual differences in emotion regulation moderate the associations between empathy and affective distress. Motivation and Emotion, 42: 602–613.

Preston, S. D., & de Waal, F. B. M. (2001) Empathy: Its ultimate and proximate bases. Behavioural & Brain Sciences, 25: 1–20.

Ray, R. D., McRae, K., Ochsner, K. N., & Gross, J. J. (2010). Cognitive reappraisal of negative affect: converging evidence from EMG and self-report. Emotion, 10: 587–92.

Reniers, R. L., Corcoran, R., Drake, R., Shryane, N. M., & Vollm, B. A. (2011). The QCAE: a Questionnaire of Cognitive and Affective Empathy. Journal of Personality Assessment, 93: 84– 95.

Rueckert, L., Branch, B., & Doan, T. (2011). Are Gender Differences in Empathy Due to Differences in Emotional Reactivity? Psychology, 2: 574–578.

Sabbagh, M. A., Xu, F., Carlson, S. M., Moses, L. J., & Lee, K. (2006). The Development of Executive Functioning and Theory of Mind: A Comparison of Chinese and U.S. Preschoolers. Psychological Science, 17: 74–81.

Samson, A. C., Huber, O., & Gross, J. J. (2012). Emotion regulation in Asperger’s syndrome and high-functioning autism. Emotion, 12: 659.

Sheppes, G., & Gross, J. J. (2011). Is timing everything? Temporal considerations in emotion regulation. Personality & Social Psychology Review, 15: 319–31.

Singer, T., & Lamm, C. (2009). The social neuroscience of empathy. Annals of the New York Academy of Sciences, 1156: 81–96.

Sonnby-Borgstrom, M. (2002). Automatic mimicry reactions as related to differences in emotional empathy. Scandinavian Journal of Psychology, 43: 433–43.

Strayer, J. (1993). Children’s Concordant Emotions and Cognitions in Response to Observed Emotions. Child Development, 64: 188–201.

Thompson, R. A. (2011). Emotion and Emotion Regulation: Two Sides of the Developing Coin. Emotion Review, 3: 53.

Tottenham, N., Hare, T. A., & Casey, B. J. (2011). Behavioral assessment of emotion discrimination, emotion regulation, and cognitive control in childhood, adolescence, and adulthood. Frontiers in Psychology, 16: 2–39.

Tully, E. C., Ames, A. M., Garcia, S. E., & Donohue, M. R. (2016). Quadratic Associations Between Empathy and Depression as Moderated by Emotion Dysregulation. Journal of Psychology, 150: 15–35.

Urry H. L., van Reekum, C. M., Johnstone, T., & Davidson, R. J. (2009). Individual differences in some (but not all) medial prefrontal regions reflect cognitive demand while regulating unpleasant emotion. NeuroImage. 47:852–863.

Wang, H. Y., Xu, G. Q., Ni, M. F., Zhang, C. H., Sun, X. P., Chang, Y., & Zhang, B. W. (2017). Neural mechanisms of implicit cognitive reappraisal: Preceding descriptions alter emotional response to unpleasant images. Neuroscience, 347: 65–75.

Wild, B., Erb, M., & Bartels, M. (2001). Are emotions contagious? Evoked emotions while viewing emotionally expressive faces: quality, quantity, time course and gender differences. Psychiatry Research, 102: 109–124.

Wu, L., Winkler, M. H., Andreatta, M., Hajcak, G., & Pauli, P. (2012). Appraisal frames of pleasant and unpleasant pictures alter emotional responses as reflected in self-report and facial electromyographic activity. International Journal of Psychophysiology, 85: 224–229.

Zaki, J. (2014). Empathy: a motivated account. Psychological Bulletin, 140: 1608–47.

