## Supplemental Materials for "How Cognitive and Affective Empathy Relate to Emotion Regulation: Divergent Patterns from Trait and Task-Based Measures"

Supplementary Table S1

*Spearman's Correlation Coefficients and p-Values for Study 1 Correlations between QCAE and ERQ-Reappraisal Following Removal of Univariate and Bivariate Outlier Cases*

| | ERQ-Reappraisal |
|---|---|
| QCAE-Cognitive | *rho*(202) = .22, *p* = .001 |
| QCAE-Affective | *rho*(205) = .07, *p* = .33 |

Supplementary Table S2

*Negative IAPS Image Numbers and Corresponding Neutralising and Descriptive Frame Sentences from the Study 2 Implicit Reappraisal Task*

| IAPS Number | Neutralising Frame Sentence | Descriptive Frame Sentence |
|---|---|---|
| 2205 | The man's wife was ill but is fully recovering | The man holds his sick wife's hand |
| 2700 | These women are overwhelmed with joy at a wedding | The women are crying in a group |
| 3030 | With medical care he will only have a small scar | He will have scarring from the flesh wound |
| 3051 | The amount of blood makes it look worse than it is | She has some cuts and bruises on her face |
| 3101 | With good medical care, this man's burns will heal well | This man has facial burns which require medical care |
| 3180 | This is an actress in a film about domestic abuse | She has been beaten and has a black eye |
| 3300 | Thanks to medical care she will recover from her illness | She is receiving medical care in a hospital |
| 3350 | Thanks to early care this baby develops into a healthy toddler | While in early care this premature baby is being monitored |
| 3500 | The man escapes unharmed | The man is being held at gun-point |
| 6212 | This soldier is protecting the child | The soldier is watching the child run |
| 6350 | This is an actress in a self-defence training video | The man is holding a knife to the woman's throat |
| 6834 | Mother and child are protected by the policemen's actions | The police are arresting a man in front of a mother and child |
| 6838 | The child is scared but will come to no harm | The child is crying in front of the car |
| 9041 | She is just a little shy | She is hiding her face |
| 9220 | The couple can take comfort from one another | The couple are at the graveside |
| 9250 | These doctors will save the woman's life | The medics are taking the young woman to medical care |
| 9429 | The children are being protected by their mothers | The children and mothers are running down the road |
| 9561 | The kitten can be treated by a vet | The kitten has an unhealthy-looking eye |
| 9900 | Nobody was seriously injured in this car accident | A large number of firefighters were sent to the scene |
| 9921 | The firefighters get this woman to safety just in time | The firefighters are carrying a woman from a burning building |

Supplementary Table S3

*Neutral IAPS Image Numbers and Corresponding Descriptive Frame Sentences from the Study 2 Implicit Reappraisal Task*

| IAPS Number | Descriptive Frame Sentence |
|---|---|
| 2025 | She is holding her finger to her mouth |
| 2038 | The woman is reading something on her laptop |
| 2102 | The man reads the stock report every morning |
| 2280 | The boy has brown eyes and short hair |
| 2385 | Her hair is covering her face |
| 2393 | These workers are checking the settings of a complicated machine |
| 2394 | The doctor is checking a patient’s measurement |
| 2396 | The couple are walking down a flight of stairs holding a cat basket |
| 2441 | She has blue eyes and long hair |
| 2480 | The man is holding the curtains to look out the window |
| 2491 | He is checking his temperature with a thermometer |
| 2500 | The man has a large white beard |
| 2515 | The woman and children are picking strawberries |
| 2579 | The chefs are making dumplings in the market |
| 2580 | These men play chess three times a week |
| 2593 | This café has outdoor seating |
| 2597 | People are queuing to buy tuna at the fish market |
| 2635 | The cowboy is cold in the snow holding his coat |
| 2749 | The man is smoking a cigar and drinking wine in his house |
| 7550 | The man is working on an old engineering program |

Supplementary Table S4

*Spearman's Correlation Coefficients and p-Values for Study 2 Correlations between QCAE and Implicit Reappraisal Task Metric Following Removal of Univariate and Bivariate Outlier Cases*

| | Implicit Reappraisal |
|---|---|
| QCAE-Cognitive | *r*(83) = -.097, *p* = .38 |
| QCAE-Affective | *rho*(83) = .44, *p* < .001 |
